# Haplotype-phased assemblies of the two poplar rust fungi species: *Melampsora larici-populina* and *Melampsora allii-populina*

**DOI:** 10.64898/2026.04.23.720444

**Authors:** Emma Corre, Emmanuelle Morin, Jana Sperschneider, Ammar Abdalrahem, Michael Pernaci, Igor V. Grigoriev, Pascal Frey, Sébastien Duplessis, Cécile Lorrain

**Affiliations:** Université de Lorraine, INRAE, UMR 1136 IAM, F-54000, Nancy, France; CSIRO Agriculture and Food, Canberra, ACT 2601, Australia; US Department of Energy Joint Genome Institute, Lawrence Berkeley National Laboratory, Berkeley, California, USA; Department of Plant and Microbial Biology, University of California Berkeley, Berkeley, California, USA; Institute of Integrative Biology, ETH Zürich, Zürich, Switzerland

**Keywords:** Poplar rust, PacBio HiFi sequencing, *Melampsora* spp., haplotype-resolved assemblies, Pucciniales, transposable elements

## Abstract

Dikaryotic rust fungi maintain two distinct haploid nuclei for most of their life cycle, making their large, repeat-rich genomes difficult to assemble and phase. Here we present haplotype-phased, near chromosome-scale genome assemblies for the poplar rust pathogens *Melampsora larici-populina* 98AG31 and *Melampsora allii-populina* 12AY07, generated using PacBio HiFi sequencing and Hi-C-guided scaffolding. For each species, we resolved 18 chromosomes per haplotype, providing the first near chromosome-level representations of poplar rust fungal species. *M. larici-populina* diploid assembly spans ~203 Mb, while *M. allii-populina* reaches ~416 Mb, with high completeness and strong collinearity between haplotypes. Compared with previous fragmented or collapsed references, these assemblies greatly improve contiguity, recover centromeric and telomeric features, and support the transposable element-driven genome size expansion in *M. allii-populina*. The haplotype-aware annotations of genes and predicted effectors derived from these resources will enable detailed analyses of genome architecture, repeat dynamics, and key loci such as avirulence genes. Together, these assemblies provide a robust genomic resource for investigating host adaptation, virulence evolution, and population diversity in poplar rust fungi.

## INTRODUCTION

Rust fungi (or Pucciniales) have remarkable biological and genomic complexity, particularly in their life cycles and nuclear organization (Henningsen et al., 2025). The Pucciniales represent the largest order of obligate biotrophic fungal pathogens, with >8,000 described species (Aime & McTaggart, 2021), infecting diverse hosts, including major crops such as wheat, soybean, and oat (Henningsen et al., 2025). Many rust fungal species follow a heteroecious and macrocyclic life cycle, alternating between two taxonomically distinct host plants and producing multiple spore types (Lorrain et al., 2019). In addition, rust fungi maintain a dikaryotic stage for most of their life cycle, with two haploid nuclei coexisting in the same spore but remaining physically separated (Henningsen et al., 2025). Unlike diploid organisms, where homologous chromosomes co-exist in a single nucleus, rust fungal genomes display distinct haplotypes in these two nuclei. This unique genomic arrangement, along with a high repeat content, has historically complicated efforts to generate fully phased telomere-to-telomere (T2T) genome assemblies (Duplessis et al., 2011). Long-read sequencing spanning repetitive regions supports improvement in contiguity, while Hi-C contact maps support chromosome-scale scaffolding, haplotype phasing, and structural genome resolution (Edwards et al., 2022; Henningsen et al., 2022, 2024; Li et al., 2019; Liang et al., 2023; Sperschneider et al., 2025; Tobias et al., 2021; Wang et al., 2024; Wu et al., 2021; Xia et al., 2022). These phased references enable fine-scale analyses of structural variation and TE landscapes, and facilitate the discovery of features linked to host adaptation (Corre et al., 2025; Henningsen et al., 2024; Li et al., 2019).

The poplar rust fungi *Melampsora larici-populina* and *Melampsora allii-populina* are damaging pathogens of poplar with major implications for forestry, bioenergy production, and ecosystem stability (Frey et al., 2005). Both infect poplar during the telial stage but differ in their aecial hosts for sexual reproduction, alternating on larch (*Larix* spp.) and Alliaceae species, respectively (Hacquard et al., 2011; Vialle et al., 2011). Early genome work on *M. larici-populina* revealed expansions in effector gene families, genome plasticity, and coordinated expression of candidate secreted effector proteins across development and host interaction (Duplessis et al., 2011; Lorrain et al., 2018). A high-density genetic map further informed recombination patterns and supported scaffolding into 18 linkage groups (Pernaci et al., 2014). This ~110 Mb reference genome represents a collapsed assembly of the two haplotypes, limiting resolution of inter-haplotype diversity and structural variants. For *M. allii-populina*, a 335.7 Mb draft assembly was generated with long reads as part of the Joint Genome Institute 1000 Fungal Genomes project (Grigoriev et al., 2014), but it remained fragmented and unphased (Duplessis et al., unpublished).

Here, we present the first haplotype-phased, near chromosome-scale genome assemblies for *M. larici-populina* and *M. allii-populina*, generated by combining PacBio long-read sequencing and Hi-C scaffolding. These references resolve both nuclear genomes of the dikaryotic state and enable comprehensive haplotype-aware gene and TE annotations, providing a foundation for comparative and functional genomics of poplar rust pathogens and for dissecting the genomic bases of biotrophic adaptation.

## MATERIALS & METHODS

### Fungal material

Urediniospores of *M. larici-populina* strain 98AG31 and *M. allii-populina* strain 12AY07 were multiplied and collected from poplar leaves as previously described in Petre et al., (2012). *In planta* and spore material used for RNA extraction and sequencing used for gene prediction were performed as described in Lorrain et al., (2018) and Petre et al., (2012).

### PacBio HiFi DNA and Hi-C sequencing

High molecular weight DNA extractions were performed from urediniospores for *M. larici-populina* 98AG31 and *M. allii-populina* 12AY07, as described in (Persoons et al., 2014). Briefly, 50-100 mg of urediniospores were ground with liquid nitrogen-cooled mortar and pestle, and DNA was extracted using a CTAB buffer followed by phenol-chloroform purification (Persoons et al., 2014). RNA was digested with RNaseA, and DNA was precipitated with isopropanol, washed with ethanol, and resuspended in Tris-EDTA. The resulting purified DNA samples were sent for library preparation and sequencing using the PacBio Revio platform (Novogene Europe, Germany).

Urediniospores of *M. larici-populina* 98AG31 and *M. allii-populina* 12AY07 were prepared for Hi-C analysis using the PhaseGenomics protocol (Seattle, Washington, USA). The samples were crosslinked with formaldehyde according to the manufacturer’s instructions to preserve chromatin structure. Hi-C library preparation was carried out using the Proximo Hi-C Kit (Fungal), which incorporates a four-enzyme digestion strategy specifically optimized for fungal genomes with high AT content by PhaseGenomics (Seattle, Washington, USA). The resulting libraries were sequenced on an Illumina NovaSeq X platform using paired-end sequencing to generate 50M of 75 bp paired-end reads.

### Genome assembly and scaffolding

Genome assemblies for *M. larici-populina* 98AG31 and *M. allii-populina* 12AY07 were assembled using the procedure described in Sperschneider et al., (2023). HiFi reads were assembled with hifiasm v0.19.9-r616 (Cheng et al., 2021) with Hi-C integration mode and default parameters (25.3 Gb HiFi reads and 65M Hi-C reads for *M. larici-populina* 98AG31; 39.7Gb HiFi reads and 55M Hi-C reads for *M. allii-populina* 12AY07). We estimated contig coverage by re-aligning HiFi reads against the diploid assemblies using minimap2 (v2.24) (H. Li, 2021), and extracted contig coverage using bbmap package (v 38.18)(Bushnell, 2014). Contigs with less than 7X and 2X coverage were filtered from *M. larici-populina* 98AG31 and *M. allii-populina* 12AY07 assemblies, respectively. The two haplotype assemblies from hifiasm were combined into a diploid genome assembly, and mitochondrial and contaminant contigs were removed based on BLAST similarity searches (v2.12)(Altschul et al., 1990). Sequences with BLAST hits assigned to taxa outside the Pucciniales order were also removed.

Mitochondrial contigs were identified based on high coverage and low GC content, then confirmed via BLASTn against the NCBI RefSeq mitochondrial genome database (≥90% identity, with DUST masking parameters; https://github.com/JanaSperschneider/GenomeAssemblyTools/). For each haplotype, the most represented contigs ranging from 70 to 90 kb in length were extracted as putative mitochondrial contigs. We assessed the completeness of each draft assembly using BUSCO v5 (Simão et al., 2015).

To phase the dikaryotic genome assemblies, we used NuclearPhaser (Duan et al., 2022), a graph-based pipeline that assigns contigs to haplotypes based on Hi-C chromatin contact data and gene homology (Duan et al., 2022). Briefly, NuclearPhaser identifies homologous contigs using gene and BUSCO duplication patterns to build scaffold bins, then uses Hi-C contact graphs to assign these bins to haplotypes through community detection. Contigs with inconsistent Hi-C signals are flagged as potential phase switches, which can be manually corrected to improve phasing accuracy. We generated gene hit tables using Biokanga v4.4.2 (Whan et al., 2019) and BUSCO hit tables using BUSCO v5 (Simão et al., 2015). Hi-C reads were mapped to each diploid assembly of *M. larici-populina* 98AG31 and *M. allii-populina* 12AY07 using bwa-mem2 v 0.7.17 (Vasimuddin et al., 2019). Hi-C interaction matrices were constructed using the *hicBuildMatrix* function from the HiCExplorer (Ramírez et al., 2018) suite with a 100 kb bin size and corrected with iterative correction and eigenvector decomposition (ICE) via *hicCorrectMatrix*. The HiCExplorer matrices were converted into *ginteractions* format using *hicConvertFormat*. Phase switches were corrected before running the NuclearPhase pipeline for a second round to validate the phase switches and scaffolding the assemblies. For *M. larici-populina* 98AG31, the scaffolding of the fully phased assembly was finalized using Ragtag v2.1.0 (Alonge et al., 2022) *scaffold* function with default parameters, using the *M. larici-populina* 98AG31 version 2 as reference (Persoons et al., 2022). The *M. larici-populina* 98AG31 genome version 2 was generated at JGI from Sanger sequences of genome assembly version 1 (Duplessis et al., 2011), reassembled with ARACHNE (Batzoglou et al., 2002) and anchored to a genetic map built from resequencing data of a selfing progeny of *M. larici-populina* 98AG31 (Pernaci et al., 2014). Scaffolds were re-ordered following *M. lini haplotype C* haplotype-phased assembly (Sperschneider et al., 2025). For *M. allii-populina* 12AY07 scaffolding, we constructed a *de novo* genome diploid assembly using the assembly from the first round of NuclearPhaser with the pipeline HapHiC (Zeng et al., 2024). For this, we re-mapped the Hi-C reads against the phased switch NuclearPhaser assembly with bwa-mem2 and ran the HapHiC pipeline with default parameters and expected number of chromosomes set at 36. We then compared the resulting HapHic diploid assembly to our previous assembly to separate the two haploid genomes with whole-genome alignments dot plots generated with D-genies (Cabanettes & Klopp, 2018). We used alignments with *M. lini haplotype C* haplotype-phased assembly to re-order scaffolds of *M. allii-populina*. We corrected the remaining mis-assemblies using whole-genome alignments between the two haplotypes and used Ragtag *correct* to break mis-assemblies, followed by Ragtag *scaffold* to correct them (Alonge et al., 2022). Finally, we verified potential remaining misplaced contigs by running NuclearPhaser again to correct the last phase switch with the new Hi-C contact map.

We searched for telomeric repeats on contigs with FindTelomeres.py (https://github.com/JanaSperschneider/FindTelomeres). We detected the putative centromeres by visual inspection of the Hi-C contact maps for each haplotype using the HiCExplorer suite (Ramírez et al., 2018). The step-by-step workflow for *M. larici-populina* 98AG31 and *M. allii-populina* 12AY07 assemblies is available here: https://github.com/Raistrawby/MrT2T.

### Genome alignments and synteny visualization

To assess structural collinearity and divergence between haplotypes, we performed pairwise whole-genome alignments using Minimap2 v2.1 (Li, 2021) with the -x asm5 parameter, which is optimized for aligning closely related assemblies. Alignments were generated between haplotype 0 and haplotype 1 for each species. The resulting alignments were output in PAF format and processed using the paftools.js stat utility from Minimap2 to extract summary statistics, including the percentage of aligned bases between haplotypes. For synteny visualization, we used D-GENIES (Cabanettes & Klopp, 2018) to generate dot plots. Dot plots were used to visually inspect the overall genome collinearity and detect large-scale structural rearrangements.

### Detection of centromeres using Hi-C contact matrices

Putative centromeres were detected from Hi-C contact matrices. First, we generated a Hi-C contact matrix for each genome as described above. For centromere detection, we estimated the intra- and inter-chromosomal contacts across all scaffolds using a 250kb sliding-window approach. We then compared ratios between the local inter-chromosomal contacts to the average inter-chromosomal contacts per scaffold to detect the region with a peak of inter-chromosomal contacts value corresponding to the putative centromere positions (Supplementary Table 1). Finally, each centromere was classified as: validated, potentially close and not detected, depending on the manual confirmation. We verified and manually corrected each predicted centromere position by visual inspections of inter-chromosomal Hi-C contact maps. All scripts for the analyses and visualization are available at https://github.com/Raistrawby/MrT2T.

**Table 1:**
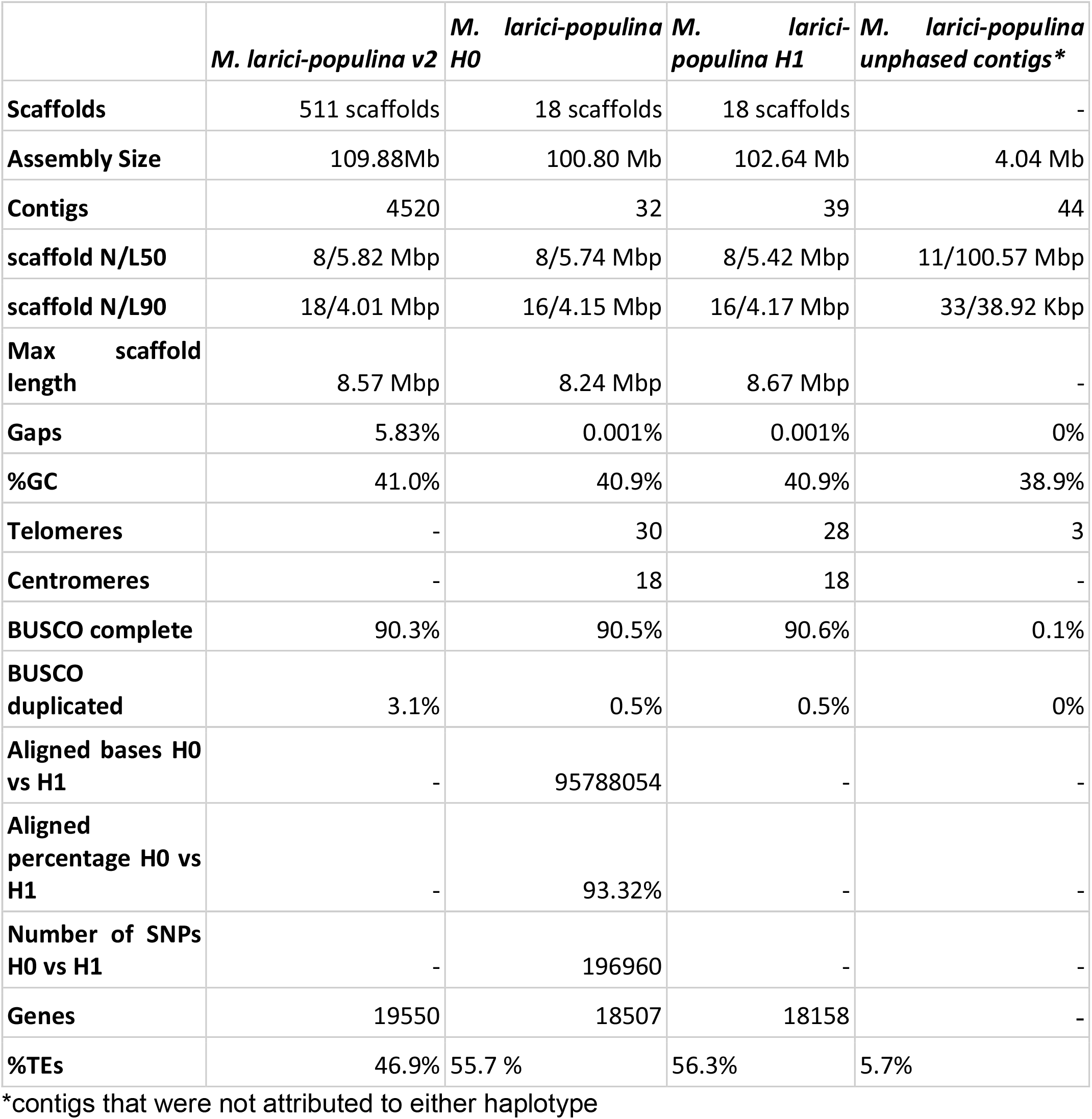
Assembly metrics of *Melampsora larici-populina* genome version 3.

### Gene and transposable elements annotations

The annotation of each haplotype for *M. larici-populina* and *M. allii-populina* was carried out using the Joint Genome Institute (JGI) Annotation Pipeline (Grigoriev et al., 2014). Briefly, the workflow identifies and masks repetitive sequences and transposable elements. Gene prediction integrates evidence from multiple sources, including ab initio, homology- and transcriptomic-based methods, to ensure accurate structural modelling. For each genomic locus, the most reliable gene model is selected to produce a high-confidence, non-redundant gene catalogue. Predicted genes were grouped into gene families and functional annotations were predicted with the JGI tools suite. Effectors were predicted with EffectorPv3.0-fungi (Sperschneider & Dodds, 2022). Mating-type genes were detected through similarity-based searches with BLASTp using *M. larici-populina* v2 annotations. Subsequently, a detailed TE classification was performed using REPETv3.0 (Flutre et al., 2011) and a curated TE library, as described in Corre et al. (2025). Gene and TE annotations and sequences are available: [zenodo].

## RESULTS AND DISCUSSION

### Complete assemblies for *M. larici-populina* and *M. allii-populina*

We generated high-quality, haplotype-phased, near chromosome-scale genome assemblies for *M. larici-populina* 98AG31 and for *M. allii-populina* 12AY07 using long-read sequencing and Hi-C scaffolding. For *M. larici-populina*, the new assembly comprises 18 chromosomes per haplotype, each representing a near-complete nuclear genome, with a total assembly size of 202.72 Mb (100.88 Mb for H0 and 102.64 Mb for H1; Table 1). The N50 for each haplotype assembly is 5.74-5.42 Mb, similar to the previous version genome, which consisted of 511 scaffolds including 18 linkage groups and an N50 of 5.82 Mb (Table 1; Figure 1). The improved assembly also closed numerous gaps compared to the previous version, with 5.83% of gaps in *M. larici-populina* 98AG31 version 2 and 001% gaps in *M. larici-populina* 98AG31 version 3 H0 and H1, respectively (Table 1; Figure 1). We mapped the centromeric regions for all four haplotypes, based on interchromosomal Hi-C contact ratios. We could annotate 18 and 16 centromeres in *M. larici-populina* haplotype 0 and haplotype 1, respectively using the inter-chromosomal contact ratios (Supplementary Table S1; Figure 2). The remaining centromeres were recovered by visual inspection of the contact maps. Finally, we detected a total of 61 telomeres, indicating that 24 scaffolds were assembled at the chromosome level (11 for H0 and 13 for H1) and 31 show at least one telomere (Table 1).

**Figure 1.**
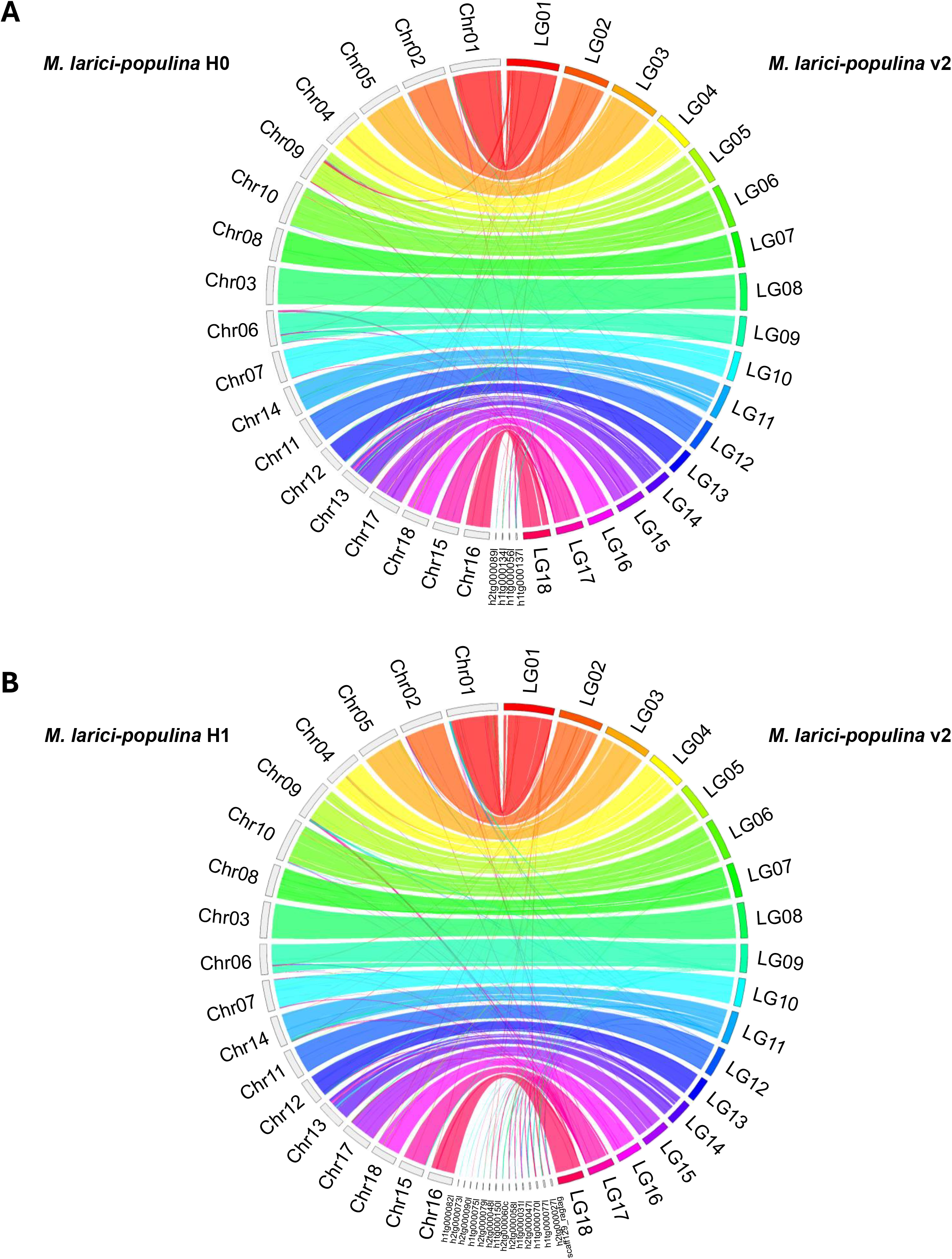
Comparison of *Melampsora larici-populina* 98AG31 genome assemblies. Circos plots showing synteny between the *M. larici-populina* v2 reference assembly and A) the *M. larici-populina* v3 haplotype 0 assembly and B) the *M. larici-populina* v3 haplotype 1 assembly. The plots were generated using Circos with the following parameters: window size = 50 kb, minimum gap size = 5 kb, minimum quality score = 20, scaffold break threshold = 50 kb gap, and a minimum link size = 10 kb (Krzywinski et al., 2009).

**Figure 2.**
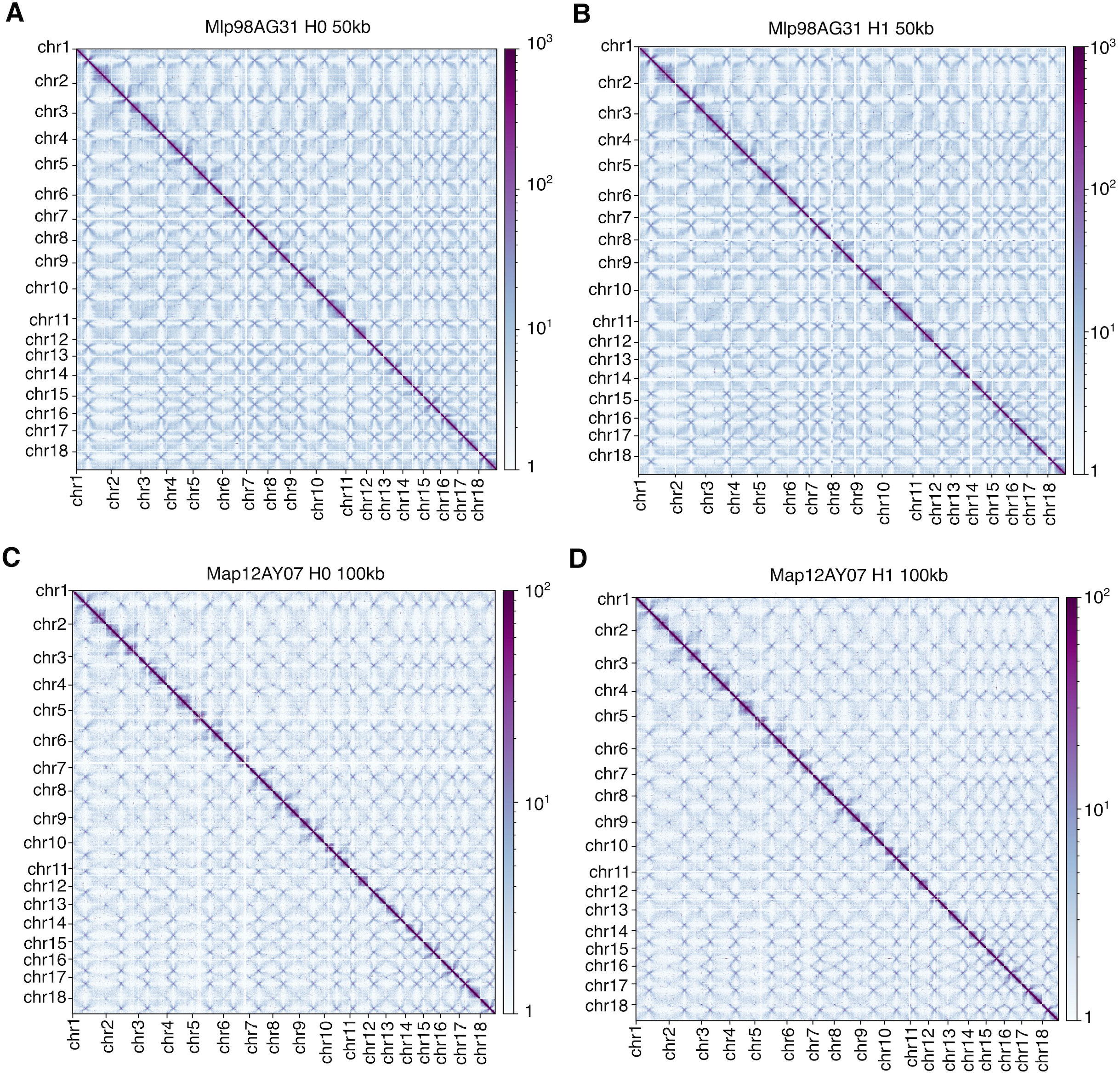
Genome-wide Hi-C contact maps. Heatmaps of normalized Hi-C contact frequency at 50 kb and 100 kb bin resolution for: A) *M. larici-populina* 98AG31 haplotype 0, B) *M. larici-populina* 98AG31 haplotype 1, C) *M. allii-populina* 12AY07 haplotype 0, D) *M. allii-populina* 12AY07 haplotype 1. The dark purple diagonal indicates intra-chromosomal interactions, while the bowtie-shaped crosses correspond to centromeric regions.

*M. allii-populina* 12AY07 exhibited a diploid genome size of 415.95 Mb with 224.03-219.39 Mb per haplotype and 8.53 Mb of unplaced contigs (Table 2). In contrast, the previous version 1 assembly for *M. allii-populina* was fragmented, containing 3637 scaffolds over 335.73Mb. The N50 for each haplotype assembly is 13.31-12.91 Mb for H0 and H1, respectively, representing an increase in contiguity compared to the version 1 genome, which consisted of an N50 of only 0.19 Mb (Table 2). For *M. allii-populina*, we automatically annotated 14 centromere coordinates for each haplotype using the inter-chromosomal contact ratios, due to the lower resolution of the Hi-C matrices for this species. However, as for *M. larici-populina*, 18 centromeres for both haplotypes were detected using visual inspection (Supplementary Table S1; Figure 2). At least one telomeric repeat was identified for 17 scaffold ends in H0 and H1, each, and we annotated both telomere for 9 scaffolds in H0 and 6 in H1, supporting the near T2T level of this assembly.

**Table 2:** Assembly metrics of *Melampsora allii-populina* genome version 2. *contigs that were not attributed to either haplotype

|  | *M. allii-populina*<br>v1 | *M. allii-populina*<br>H0 | *M. allii-populina*<br>H1 | *M. allii-populina*<br>unphased contigs* |
| --- | --- | --- | --- | --- |
| **Scaffolds** | 3637 scaffolds | 18 scaffolds | 18 scaffolds | - |
| **Assembly Size** | 335.73Mb | 224.03Mb | 219.39Mb | 8.53 Mb |
| **Contigs** | 3637 | 33 | 46 | 116 |
| **scaffold N/L50** | 440/191.40 Kbp | 8/13.31 Mbp | 8/12.91 Mbp | 30/79.11 Kbp |
| **scaffold N/L90** | 2188/35.5 Kbp | 16/9.29 Mbp | 16/9.11 Mbp | 92/41.98 Kbp |
| **Max scaffold length** | 1.29 Mbp | 17.52 Mbp | 16.99 Mbp | 312.62 Kbp |
| **Gaps** | 0% | 0% | 0.001% | 0% |
| **%GC** | 40.0% | 40.0% | 40.0% | 38% |
| **Telomeres** | - | 26 | 24 | 2 |
| **Centromeres** | - | 18 | 18 | - |
| **BUSCO complete** | 89.7% | 90.7% | 86.1% | 0.3% |
| **BUSCO duplicated** | 22.1% | 1.1% | 1.1% | 0.1% |
| **Aligned bases H0 vs H1** | - | 192817739 | - | - |
| **Aligned percentage H0 vs H1** | - | 87.9% | - | - |
| **Number of SNPs H0 vs H1** | - | 906156 | - | - |
| **Genes** | 23089 | 17745 | 17494 | - |
| **%TEs** | 74.8% | 82.3% | 82.0% | 2.7% |

Both assemblies exhibit high completeness, with BUSCO scores of 90.5% and 88.4% for *M. larici-populina* and *M. allii-populina*, respectively, based on the Basidiomycota reference set (Tables 1 and 2). In addition, we found 0.5% and 1.1% of BUSCO duplicated levels in *M. larici-populina* v3 and *M. allii-populina* v2, compared to 3.1% and 22.1% in the previous versions of *M. larici-populina* v2 and *M. allii-populina* v1, respectively. This reduction of duplication levels supports the haplotype resolution of these new assemblies. We have generated two fully phased and near T2T reference genome assemblies for each of of two poplar rust species. These phased, near-T2T assemblies provide a critical foundation for resolving genome structure, functional gene repertoires, and the evolutionary dynamics of repeats in obligate biotrophs with dikaryotic life stages. In rust fungi, haplotype resolution is particularly important to quantify haplotypic divergence, detect structural rearrangements, and accurately reconstruct key loci such as mating-type regions. The haploid genome sizes recovered here are consistent with previous genome size estimates and comparative analyses within Pucciniales (Corre et al., 2025; Duplessis et al., 2011; Sperschneider et al., 2025). The strong improvements relative to earlier assemblies, especially for *M. allii-populina*, underscore the effectiveness of integrating long-read sequencing with Hi-C–based phasing for large, repeat-rich, heterozygous rust genomes (Henningsen et al., 2022). This framework has proven robust across Pucciniales, including *M. lini* and other rusts (Henningsen et al., 2025; Sperschneider et al., 2025), and our results further support its general applicability. Finally, both assemblies resolve 18 chromosomes per haplotype with identifiable telomeric and centromeric features, consistent with the conserved karyotype (n = 18) reported for most rust fungi (Henningsen et al., 2025). Together, these resources substantially advance *Melampsora* genomics and provide high-resolution references for investigating haplotype evolution, genome architecture, and adaptive processes in poplar rust pathogens.

### The dikaryotic genomes of *M. larici-populina* 98AG31 and *M. allii-populina* 12AY07 display similar levels of heterozygosity and strong inter-haplotype collinearity

The dikaryotic genomes of *M. larici-populina* 98AG31 and *M. allii-populina* 12AY07 exhibit both high heterozygosity and strong collinearity between haplotypes, consistent with the long-term maintenance of divergent nuclei in rust fungi. We assessed the heterozygosity levels of raw reads using a 21-mer-based approach. Heterozygosity was estimated at 1.02% for *M. larici-populina* and 1.01% for *M. allii-populina* (Figure 3), indicating substantial allelic divergence between the two dikaryotic haplotypes. This is consistent with sustained co-existence of divergent nuclei in dikaryotic rusts and suggests that haplotypic divergence is maintained genome-wide, as observed in other Pucciniales species (Schwessinger et al., 2018; Wang et al., 2024).

**Figure 3.**
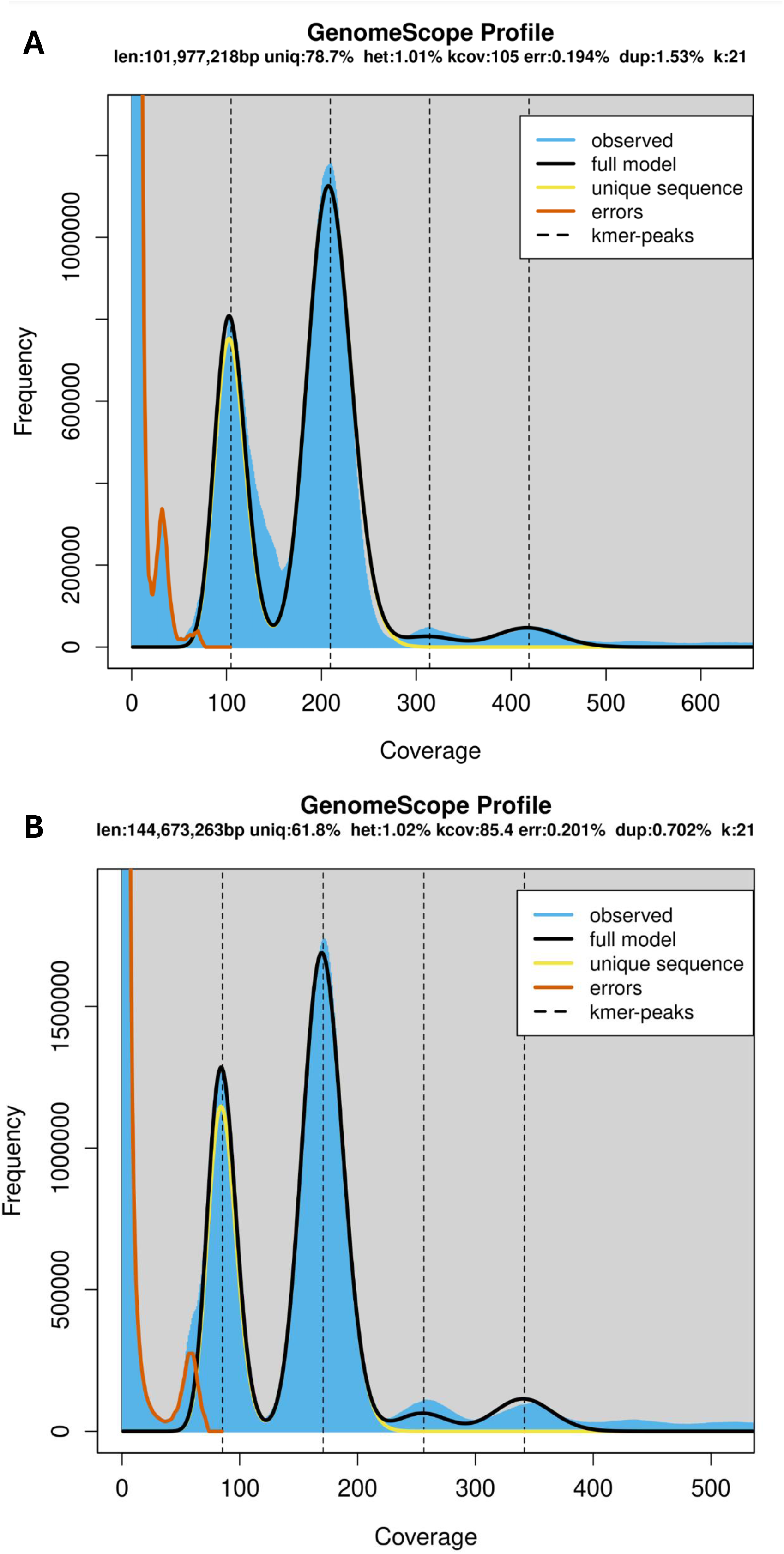
Genome size and heterozygosity estimation from HiFi read k-mer spectra. GenomeScope (Ranallo-Benavidez et al., 2020) 21-mer profiles of HiFi sequencing reads from *M. larici-populina* 98AG31 with an estimated genome heterozygosity (het) of 1.01%, and *M. allii-populina* 12AY07 with an estimated genome heterozygosity of 1.02%. The distinct 1:2 ratio of the k-mer peaks supports the diploid nature of both species.

To evaluate large-scale genome conservation and detect potential structural rearrangements, we compared haplotypes using whole-genome alignments visualized with collinearity dot plots (Supplementary Figures S1 and S2). In *M. larici-populina*, 93.3% of the haplotype 0 sequence aligned to haplotype 1, reflecting strong inter-haplotype conservation (Table 1). In *M. allii-populina*, the aligned fraction was lower (87.2%; Table 2), consistent with increased structural divergence relative to *M. larici-populina*. Nevertheless, both species exhibited high overall collinearity between haplotypes, with no evidence for large-scale chromosomal rearrangements (Supplementary Figures S1C and S2C). In both species, we identified a single major non-syntenic region (~1.5-2 Mb) on chromosome 9, characterized by little to no detectable alignment between haplotypes (Supplementary Figure S1D, Supplementary Figure S2D). This region corresponds to the mating-type locus and includes the pheromone receptor (PR/STE) genes (Supplementary Table S2), consistent with observations in other Pucciniales, including *Puccinia graminis* f. sp. *tritici, Puccinia striiformis* f. sp. *tritici* and *M. lini*, where mating-type loci occur within divergent, haplotype-specific genomic regions (Cuomo et al., 2016; Sperschneider et al., 2025; Wang et al., 2024). Such localized divergence was suggested to reflect recombination suppression around mating-type loci, which may contribute to the long-term maintenance of mating compatibility determinants and inter-haplotype differentiation (Sperschneider et al., 2025). Together, our results indicate that both poplar rust genomes are characterized by extensive synteny between haplotypes, punctuated by a single large PR genes-associated divergent region, thereby reinforcing an emerging model of conserved karyotype organization and mating-type architecture across rust fungi. Whether such divergence reflects functional diversification of mating compatibility or structural barriers to recombination remains to be investigated. As proposed for *M. lini*, additional population-scale phased assemblies will be required to test whether mating-type structure in *M. larici-populina* and *M. allii-populina* conforms to a classic tetrapolar system or represents a more complex configuration (Lawrence, 1980).

### Improvement of the transposable element annotations in the haplotype-phased assemblies

TEs were annotated in each haplotype of the two species and classified at the Class, Order, and Superfamily (SF) levels (Supplementary Tables S3 and S4). The phased assemblies revealed differences in the overall TE genome coverage relative to previously published unphased references. The TE content in *M. larici-populina* v2 represented 47% of the genomes, while we accounted for 55.7% and 56.3% of TEs in haplotype 0 and haplotype 1 of *M. larici-populina* v3.0, respectively (Table 1, Supplementary Table S3). This indicates that the improvement in genome continuity (5% gaps in version 2 versus 0.001% gaps in the haplotype-phased assemblies) allowed for a more comprehensive recovery of repetitive elements, capturing TE sequences that were previously collapsed or fragmented in the earlier assembly. However, we still observed that *M. larici-populina* genome displays slightly more Class II DNA transposons than Class I retrotransposons, with 28.8-27.4% Class II and 20.8-22.5% Class I in haplotype 0 and 1, respectively, while we found 15.4% and 23.2% in the earlier genome version (Supplementary Table S3). Similarly, *M. allii-populina* haplotype-phased assemblies displayed 82.3-82.0% of TEs in haplotypes 0 and 1, respectively, while the previous version accounted for 74.8% of TEs (Table 2; Supplementary Table S4). Class I elements cover 49.5-49.2% of haplotypes 0 and 1, respectively, and Class II elements represent 27.8-27.9% of haplotypes 0 and 1, respectively (Supplementary Tables S3). At the Order level, LTR retrotransposons dominate the TE landscapes of *M. allii-populina* (43.5% of LTRs in haplotype 0 and 43.9% in haplotype 1), whereas *M. larici-populina* displayed a higher proportion of TIR DNA transposons (Supplementary Tables S3 and S4). In the latter, TIR elements accounted for 23.1% in haplotype 0 and 21.9% in haplotype 1, similar to the coverage found in the previous version and remain the most abundant order in the phased genome (Supplementary Tables S3 and S4). At the Superfamily level, Gypsy-elements remain the most prevalent in *M. allii-populina* (30.0% and 30.3% of the genome in haplotype 0 and 1), while in *M. larici-populina*, Gypsy elements constitute the second most abundant category after TIR elements (with ~8% in Gypsy in both haplotypes; Supplementary Tables S3 and S4). Taken together, these findings confirm that haplotype-resolved assemblies offer a significantly enhanced representation of the TE landscape in the two poplar rust species.

The increased contiguity of genome assemblies also improves annotation of TEs, which play a dominant role in shaping rust genomes (Corre et al., 2025). TE content was 25% higher in *M. larici-populina* and 10% higher in *M. allii-populina* in our haplotype-phased assemblies compared to previous versions, supporting the observation that collapsed assemblies underestimate repetitive regions (Edwards et al., 2022; Henningsen et al., 2025). Despite such differences between genome assemblies, the overall tendencies in TE composition remained consistent between assembly versions for each species, indicating that while total TE content was underestimated previously, the dominant TE classes were already correctly identified. The improved continuity and haplotype resolution in our assemblies enabled the recovery of previously collapsed or fragmented elements, resulting in more accurate estimates of repeat abundance. In *M. allii-populina*, Class I LTR retrotransposons, particularly Gypsy elements, remained the most abundant, while *M. larici-populina* continued to show an enrichment of Class II TIR DNA transposons. These refined annotations align with TE compositions reported in other rust fungi and illustrate the value of long-read, chromosome-scale assemblies in capturing complex repetitive landscapes (Sperschneider et al., 2025).

### Gene annotations in *M. larici-populina* and *M. allii-populina* haplotype-phased assemblies

Gene models were predicted independently for each haplotype of the phased assemblies of *M. larici-populina* and *M. allii-populina*, yielding comprehensive gene catalogues. In *M. larici-populina*, we annotated 18,507 (H0) and 18,158 (H1) protein-coding genes, compared with 19,550 genes reported for the prior reference assembly (v2.0)(Table 1). In *M. allii-populina*, we annotated 17,745 (H0) and 17,494 (H1) protein-coding genes, whereas the previous collapsed assembly reported 23,089 genes (Table 2). The reduced gene counts in the phased resources are consistent with improved resolution of allelic regions that can otherwise inflate gene numbers in collapsed assemblies, and they support the presence of haplotype-resolved homologous genes that are recovered only when the two haplotypes are represented separately (Sperschneider et al., 2025). Functional annotation using KOG classifications and secreted protein (SPs) predictions indicated that both species encode repertoires consistent with obligate biotrophy and plant-pathogen interaction (Duplessis et al., 2021). KOG assignments were obtained for 5,861 (H0) and 5,765 (H1) genes in *M. larici-populina* and for 6,054 (H0) and 5,923 (H1) genes in *M. allii-populina* (Supplementary Tables S5). Across both species and haplotypes, annotated genes were broadly distributed among core functional groupings, including cellular processes and signaling (1,618-1,828 genes), information storage and processing (1,249-1,297 genes), and metabolism (1,780-1,844 genes), with a substantial fraction classified as poorly characterized (1,089-1,126 genes). Within cellular processes and signaling, prominent categories included posttranslational modification/protein turnover (KOG O), signal transduction (KOG T), and intracellular trafficking/secretion (KOG U), consistent with the importance of secretory and regulatory functions in rust fungi (Supplementary Table 5) (Duplessis et al., 2021). We identified 2,110 and 2,103 predicted SPs in *M. larici-populina* H0 and H1, respectively (11.40% and 11.58% of predicted genes), and 1,838 and 1,793 in *M. allii-populina* H0 and H1, respectively (10.36% and 10.25%) (Supplementary Table S5). In addition, 1,242 (58.9% of SPs) and 1,244 (59.1% of SPs) predicted effectors were identified in *M. larici-populina* H0 and H1 secretomes, respectively, while we predicted 1,060 (57.7% of SPs) and 1,025 (57.2% of SPs) effectors in *M. allii-populina* H0 and H1, respectively (Supplementary Table S5). The detailed functional annotations for each assembly are available in Supplementary Tables S6-S9. Taken together these results highlight that, despite the larger genome size of *M. allii-populina*, its predicted gene content is not expanded relative to *M. larici-populina*, supporting that genome expansion is largely decoupled from gene number, as also noted previously and in other Pucciniales species (Corre et al., 2025).

## CODE AVAILABILITY

All custom scripts generated for data analyses and visualization are available at https://github.com/Raistrawby/MrT2T.

## DATA AVAILABILITY

Genome assemblies, Hi-C matrices, gene annotations and TE annotations are available at https://zenodo.org/uploads/19251534. Long-read sequencing reads for each species are available under the NCBI Sequence Read Archive BioProject accession ID PRJNA1453918. Hi-C sequencing reads are available under Gene Expression Omnibus repository GSE328168. RNA sequencing reads are available under Gene Expression Omnibus repository GSE106863 and under the NCBI Sequence Read Archive BioProject accession ID PRJNA1454804. Genome assemblies and annotation are also available from JGI MycoCosm portals: http://genome.jgi.doe.gov/Melap2_H0 and http://genome.jgi.doe.gov/Mellp3_H1 for *M. larici-populina v3*.*0*; http://genome.jgi.doe.gov/Melap2_H0 and http://genome.jgi.doe.gov/Melap2_H1 for *M. allii-populina v2*.*0*.

## ACKNOWLEDGMENTS

We thank Prof. Bruce A. McDonald for hosting Emma Corre during a four-months research stay, where part of the data analyses was performed. We are grateful to Jérémy Pétrowski for his excellent technical help in producing poplar rust spores for genome sequencing.

## FIGURE LEGENDS

**Supplementary Figure S1. Whole-genome dot plots of *Melampsora larici-populina* 98AG31 haplotype-phased assemblies and version 2 comparisons**. A) Synteny dot plot of the newly assembled haplotype 0 (x-axis) against the previous reference genome version of *M. larici-populina* v2 (y-axis). B) Synteny dot plot of the newly assembled haplotype 1 (x-axis) against the *M. larici-populina* v2 (y-axis). C) Synteny dot plot between haplotype 0 (x-axis) and haplotype 1 of *M. larici-populina* (y-axis), showing high collinearity with limited structural variation. C) Close-up view of scaffold 9 synteny dot plot between haplotypes 0 (x-axis) and 1 (y-axis), highlighting a ~1.5-1.7 Mb non-syntenic region. This region corresponds to the mating-type locus containing the pheromone receptor genes, previously characterized in *M. lini*.

**Supplementary Figure S2. Whole-genome dot plots of *Melampsora allii-populina* 12AY07 haplotype-phased assemblies and *Melampsora lini C* comparisons**. A) Synteny dot plot of the *M. allii-populina* haplotype 0 (x-axis) against the previous reference genome version of *M. lini C* (y-axis). B) Synteny dot plot of the *M. allii-populina* haplotype 1 (x-axis) against the *M. lini C* (y-axis). C) Synteny dot plot between *M. allii-populina* haplotype 0 (x-axis) and *M. allii-populina* haplotype 1 (y-axis), showing high collinearity with limited structural variation. C) Close-up view of scaffold 9 synteny dot plot between *M. allii-populina* haplotypes 0 (x-axis) and 1 (y-axis), highlighting a ~2 Mb non-syntenic region. This region corresponds to the mating-type locus containing the pheromone receptor genes, previously characterized in *M. lini*.

**Supplementary Table S1. Centromere annotation based on Hi-C contact maps**.

**Supplementary Table S2. Mating-type genes BLAST search results of in *Melampsora larici-populina* v3.0 and *Melampsora allii-populina* v2.0**. MAT genes retrieved with blast against the genomes of *M. larici-populina 98AG31 haplotype 0, M. larici-populina 98AG31 haplotype 1, M. allii-populina haplotype 0, M. allii-populina haplotype 1, M. lini haplotype C, M. lini haplotype H*.

**Supplementary Table S3. Transposable elements annotations in *Melampsora larici-populina* at the Class, Order and superfamily levels**

**Supplementary Table S4. Transposable elements annotations in *Melampsora allii-populina* at the Class, Order and superfamily levels**

**Supplementary Table S5. Functional annotation summaries in *Melampsora larici-populina* (Mellp) and *Melampsora allii-populina* (Melap)**.

**Supplementary Table S6. *Melampsora larici-populina* H0 gene functions**.

Protein functional annotations are summarized per predicted protein (proteinId) and linked to the corresponding gene (gene_id) and transcript (transcriptId) identifiers from the GFF3, together with the GFF3 Name field (gff_Name) and the predicted product description (gff_product). InterPro/domain annotations are reported as a concatenated list of individual hits (domain.domain_hits) and as aggregated fields listing the InterPro entry identifiers (domain.iprId) and descriptions (domain.iprDesc), as well as the source signature database (domain.domainDb) with its accession (domain.domainId) and description (domain.domainDesc). eggNOG-derived functional assignments include predicted enzyme commission numbers (eggnog.ec), KEGG orthology identifiers (eggnog.keggKo) and associated KEGG pathway/module/reaction information (eggnog.keggPathway, eggnog.keggModule, eggnog.keggReaction, eggnog.keggRclass), KEGG BRITE categories (eggnog.brite), and transporter classification codes when applicable (eggnog.keggTc), together with the supporting search statistics (eggnog.eValue, eggnog.score). Signal peptide predictions from SignalP are provided as the predicted class (signalp.prediction), the cleavage site interval (signalp.min_location, signalp.max_location) and the signal peptide probability (signalp.sp_prob).

**Supplementary Table S7. *Melampsora larici-populina* H1 gene functions**.

Protein functional annotations are summarized per predicted protein (proteinId) and linked to the corresponding gene (gene_id) and transcript (transcriptId) identifiers from the GFF3, together with the GFF3 Name field (gff_Name) and the predicted product description (gff_product). InterPro/domain annotations are reported as a concatenated list of individual hits (domain.domain_hits) and as aggregated fields listing the InterPro entry identifiers (domain.iprId) and descriptions (domain.iprDesc), as well as the source signature database (domain.domainDb) with its accession (domain.domainId) and description (domain.domainDesc). eggNOG-derived functional assignments include predicted enzyme commission numbers (eggnog.ec), KEGG orthology identifiers (eggnog.keggKo) and associated KEGG pathway/module/reaction information (eggnog.keggPathway, eggnog.keggModule, eggnog.keggReaction, eggnog.keggRclass), KEGG BRITE categories (eggnog.brite), and transporter classification codes when applicable (eggnog.keggTc), together with the supporting search statistics (eggnog.eValue, eggnog.score). Signal peptide predictions from SignalP are provided as the predicted class (signalp.prediction), the cleavage site interval (signalp.min_location, signalp.max_location) and the signal peptide probability (signalp.sp_prob).

**Supplementary Table S8. *Melampsora allii-populina* H0 gene functions**. Protein functional annotations are summarized per predicted protein (proteinId) and linked to the corresponding gene (gene_id) and transcript (transcriptId) identifiers from the GFF3, together with the GFF3 Name field (gff_Name) and the predicted product description (gff_product). InterPro/domain annotations are reported as a concatenated list of individual hits (domain.domain_hits) and as aggregated fields listing the InterPro entry identifiers (domain.iprId) and descriptions (domain.iprDesc), as well as the source signature database (domain.domainDb) with its accession (domain.domainId) and description (domain.domainDesc). eggNOG-derived functional assignments include predicted enzyme commission numbers (eggnog.ec), KEGG orthology identifiers (eggnog.keggKo) and associated KEGG pathway/module/reaction information (eggnog.keggPathway, eggnog.keggModule, eggnog.keggReaction, eggnog.keggRclass), KEGG BRITE categories (eggnog.brite), and transporter classification codes when applicable (eggnog.keggTc), together with the supporting search statistics (eggnog.eValue, eggnog.score). Signal peptide predictions from SignalP are provided as the predicted class (signalp.prediction), the cleavage site interval (signalp.min_location, signalp.max_location) and the signal peptide probability (signalp.sp_prob).

**Supplementary Table S9. *Melampsora allii-populina* H1 gene functions**. Protein functional annotations are summarized per predicted protein (proteinId) and linked to the corresponding gene (gene_id) and transcript (transcriptId) identifiers from the GFF3, together with the GFF3 Name field (gff_Name) and the predicted product description (gff_product). InterPro/domain annotations are reported as a concatenated list of individual hits (domain.domain_hits) and as aggregated fields listing the InterPro entry identifiers (domain.iprId) and descriptions (domain.iprDesc), as well as the source signature database (domain.domainDb) with its accession (domain.domainId) and description (domain.domainDesc). eggNOG-derived functional assignments include predicted enzyme commission numbers (eggnog.ec), KEGG orthology identifiers (eggnog.keggKo) and associated KEGG pathway/module/reaction information (eggnog.keggPathway, eggnog.keggModule, eggnog.keggReaction, eggnog.keggRclass), KEGG BRITE categories (eggnog.brite), and transporter classification codes when applicable (eggnog.keggTc), together with the supporting search statistics (eggnog.eValue, eggnog.score). Signal peptide predictions from SignalP are provided as the predicted class (signalp.prediction), the cleavage site interval (signalp.min_location, signalp.max_location) and the signal peptide probability (signalp.sp_prob).

